# Multiplex LAMP for Simplified Monitoring *Cryptosporidium* and *Giardia* in Surface Water

**DOI:** 10.1101/2020.08.18.256826

**Authors:** Jerry E. Ongerth, Frhat M. A. Saaed

## Abstract

A detailed investigation was conducted into the application of an isothermal nucleic acid amplification procedure, LAMP, in conjunction with components of conventional procedure to determine ability to simplify water processing for monitoring *Cryptosporidium* and *Giardia* in surface water samples. Based on previous work demonstrating a high degree of sensitivity and selectivity of the LAMP procedure for detecting target organism DNA at low concentration in environmental media without significant interferences, this project sought to determine if LAMP primers for both organisms could be combined and applied to detect the target organisms collected from surface water samples after only minimal processing. Implementation consisted of optimizing previously described LAMPs for the SAM-1 gene of *Cryptosporidium* and for the EF1-α gene of *Giardia*; establishing their limits of detection on both specific DNA and on small numbers of oocysts and cysts processed by freeze-thaw cycles. Based on positive results from preliminary testing, a multiplexed LAMP for both organisms was applied. The multiplex lamp was able to detect the DNA of *Cryptosporidium* and of *Giardia* from samples seeded with replicates of 1, 2, 3 and 4 oocysts and cysts, both in clean water and in surface water processed by filtration and centrifuging to pellet followed only by freeze-thaw cycles. Positive and specific amplification was observed with potential for quantification using 25 µL reactions including 8 µL template, in RT qLAMP format by LightCycler. The procedure is significantly simpler and less time consuming presenting realistically practical procedure for improvement of surface water monitoring of *Cryptosporidium* and *Giardia*, compared to the widely used USEPA Method 1623

**Importance:** Pathogen identification by LAMP is well established as a sensitive and highly selective procedure, insensitive to interferences, permitting specific detection of target DNA in samples containing extraneous DNA. This combination of features was shown effective for identifying numbers of *Cryptosporidium* oocysts and *Giardia* cysts as few as 1 in unseparated particle assemblages subject only to freeze thaw cycles. Although LAMP has not been widely adopted for environmental monitoring, it has been previously demonstrated for *Cryptosporidium* and for *Giardia* in water and wastewater. Also, many clinical assays have demonstrated applicability to use where facilities and skills are limited. The simplified analysis sequence offers significant time and cost saving for routine monitoring of *Cryptosporidium* and *Giardia* in surface water samples.

## Introduction

Monitoring surface water sources of public water supply for *Cryptosporidium* and *Giardia* is important as a basis for public health protection and is required in many jurisdictions e.g. USEPA Surface Water Treatment Rule, USEPA, 2006; WHO, 2017; Australian Drinking Water Guidelines, NHMRC, 2011. The most common monitoring procedure is based on sample volumes typically 10 L but as large as 50 L, processed by standardized procedures, e.g. USEPA Method 1623 (USEPA, 2005); ISO15553 (IOS, 2006), that require filtration to form a particle concentrate that is then subject to immuno-magnetic separation (IMS), then completed by immuno-fluorescence microscopy (IFA) to identify and count *Cryptosporidium* oocysts and *Giardia* cysts.

The conventional procedure requires specialized equipment, is time consuming, requires highly trained and experienced technicians, and is expensive. The approach is subject to the criticism that it does not discriminate between live and dead organisms, and that it does not discriminate between the different species and subtypes of the organisms, some of which are more likely to infect humans, and some of which may be more virulent. Such features of oocysts & cysts can be identified by molecular approaches, most commonly PCR, (e.g. Monis et al, 2001; Smith et al, 2006) and for potential quantitation using RT qPCR, (e.g. Guy et al, 2003; Xiao et al, 2017).

Nearly 20 years ago a novel nucleic acid amplification procedure, termed LAMP, for Isothermal Loop-mediated Amplification, was developed, Notomi et al, 2000. The procedure exploits the ability of some polymerases, e.g. Bst, to amplify DNA by a strand displacement and extension mechanism taking place at a constant temperature in the range of 55°C-75°C, using 4 and 6 primers to provide for amplification. The procedure lends itself to visual endpoints by production of turbidity (precipitate) through pH change or dye release accompanying amplification. Amplification is rapid with endpoints reached in times ranging from 30 to 75 minutes. Due to the use of the multiple primer sets the procedure is highly selective, not subject to environmental interferences, and has been shown to be very sensitive, Ongerth and Karanis, 2009; Notomi et al, 2015. The combined features of LAMP make it inexpensive and suitable for field application where sophisticated lab facilities are limited.

In the 20 years since initial description of the method it has been developed predominantly for clinical applications (e.g. Poon et al, 2006; Curtis et al, 2008; Lee et al, 2009; Wei et al, 2020) but also for environmental applications (e.g. Thesikoe et al, 2005; Bakheit et al, 2008). Environmental LAMP procedures have been described for *Cryptosporidium* in water samples, Karanis et al 2007, and Inomata et al 2009, and for *Giardia*, Plutzer et al, 2009. Through collaboration with these authors, and recognizing the selectivity and sensitivity of the procedure, this project was developed, using the LAMP primer sets developed for *Cryptosporidium* and for *Giardia* by Karanis and Plutzer respectively, to determine the feasibility of simultaneous identification of *Cryptosporidium* and *Giardia* in surface water by a simplified sample processing scheme and multiplex LAMP application. Features of the LAMP procedure cited above suggest the attractive potential for using it to quantify specifically *Cryptosporidium* and/or *Giardia* DNA in crude pellets isolated from environmental water samples without the need for selective concentration of the oocysts and cysts, or for concentration of DNA from the digested pellet. Exploring this potential was the objective of the work described here.

## Materials and Methods

Oocysts and cysts of *Cryptosporidium* and *Giardia* were isolated from fresh feces collected from dairy calves and screened to identify positive samples as described previously (Ongerth & Stibbs, 1989). Isolation and cleanup were as described previously, (Ongerth, 1994), using Percol gradients to provide clean oocysts and cysts, stored at 4°C in filtered distilled water w/o preservative or antibiotics until use for production of DNA or as positive controls.

Surface water samples were collected from a local stream in new (unused) 20 L Cubitaners^®^. Water was processed as needed by 293 mm ⨯ 2μm track-etched polycarbonate filtration, and particles were recovered from the filter by squeegee as previously described (ibid), centrifuged to pellet for use in LAMP analysis and in conventional analysis using USEPA Method 1623, i.e., IMS-IFA-microscopy.

For all LAMP development and control requirements, DNA was extracted from the oocysts and cysts isolated as described above. Extraction was by QIAmp DNeasy (Qiagen, Valencia, CA) kits according to manufacturers instructions by the spin-column protocol. Briefly, approximately 10^7^ each of *Cryptosporidium* oocysts and of *Giardia* cysts were pelleted, 4500 g ⨯ 5 min, resuspended in 180 μL tissue lysis buffer, then subject to 10 freeze-thaw cycles (LiqN - 80°C), spot spun to re-condense, then allowed to digest overnight at 56°C with 20 μL proteinase K. After concentration, washing, and double elution, recovered DNA concentration was measured by MiniDrop (Thermo Scientific NanoDrop 2000c UV-Vis Spectrophotometer), and 10μL of each organism DNA was electrophoresed prior to aliquoting and storage at -80°C.

The LAMP assays were performed with primers a) for a 145-bp sequence of the SAM-1 gene specific to *Cryptosporidium parvum, C. hominis* and *C. meleagridis* (Bakheit et al. 2008) and for a 178-bp sequence of the EF1-α gene specific to *Giardia duodenalis* Assemblage A and B (accession no. AF069570) (Plutzer and Karanis, 2009). These LAMP primers have been used and evaluated as previously reported to detect and genotype *Cryptosporidium* and *Giardia* species (Bakheit et al. 2008; Plutzer and Karanis 2009). The LAMP reactions were performed in a 25 μl, reaction mix containing 12.5 μl of 2x reaction mix [10x BST buffer; 100 mM MgSO_4;_ 5 mM betaine; 1:10 Tween 20; and dNTP mix], 1.3 μL of SAM-1 or EF1-α primer mix (FIP, BIP, F3, B3, LB and LF), [(8.2 μL of particle free-Mill-Q water and 2 μL template) or (seeded 10.2 μL of particle free-Mill-Q water)]. Positive control samples were 12.5 μl of 2 x reaction mixes, 1.3 μL of SAM-1 or EF1-α primer mix, 2 µL *Cryptosporidium*-DNA template or *Giardia*-DNA template, 8.2 µL particle free-Mill-Q water. Negative controls were 12.5 μl of 2 x reaction mixes, 1.3 μL of SAM-1 or EF1-α primer mix, 2 µL of UV-irradiated filtered-Milli-Q water instead of 2 µL DNA template and 8.2 µL particle free-Mill-Q water to ensure appropriate and specific amplification parameters. LAMP was run by denaturing the sample DNA at 95°C for 30 second in a block-type PCR unit and then chilled on ice. Subsequently, 1 μL of Bst polymerase (8 U/reaction) was added to each reaction tube, followed by amplification at 63 °C, 60 minutes for *Cryptosporidium* samples (SAM-1) or 120 minutes for *Giardia* samples (EF1-α) and then heated at 80°C for 10 minutes for reaction termination. Then, 10 μL of the LAMP product with 2 μL loading buffer were mixed and electrophoresed in a 1.5% agarose gel, run at 80v for 80 minutes and then EB stained 30 min. for visualization.

Assessing the feasibility of simultaneous multiplex LAMP amplification for both *Cryptosporidium* and *Giardia* in water samples and potential for quantitative application proceeded as follows. First LAMP was applied independently to *Cryptosporidium* and to *Giardia* purified DNA to verify amplification, then to extinction dilutions of DNA to determine sensitivity. Next, ability to independently detect oocysts and cysts through freeze-thawed seeded samples and ability to detect small numbers i.e. 1-10, was determined. Then, surface water from a local stream was processed and seeded with low numbers of both organisms, i.e., 1-10, testing performance of the multiplex LAMP. Seeded pellet concentrates or concentrates from filtered surface water, typically < 100 μL, provided ample organism DNA after freeze-thaw and centrifuging to permit amplification using 10.2 μL of the homogenized supernatant as template. Finally, ability of multiplexed LAMP to detect both *Cryptosporidium* and *Giardia* in seeded samples using a combined reaction mix was determined, including ability to detect low numbers in seeded samples. The components of 25 μL reaction mix volume provide sufficient flexibility to allow addition of both SAM-1 and EF1-α primers with suitable increase of the dNTP mix to provide for amplification for both organisms. Reaction products were assessed by qRT LAMP using a Roche Light Cycler 480, including SYTO 13 for product detection in real time.

After completion of preliminary assays to establish LAMP performance, sensitivity and selectivity using i) purified organism DNA; ii) freeze-thawed organisms in dH_2_0; iii) freeze-thawed organisms seeded to surface water concentrates containing DNA from ambient organisms; iv) repeating steps ii and iii using multiplexed SAM-1 and EF1-α; a sequence of performance assays was conducted using a broad testing scheme, Figure 1, including a range of low-number organism seeding with replication and the same rigorous negative and positive controls carried throughout development.

**Figure 1.**
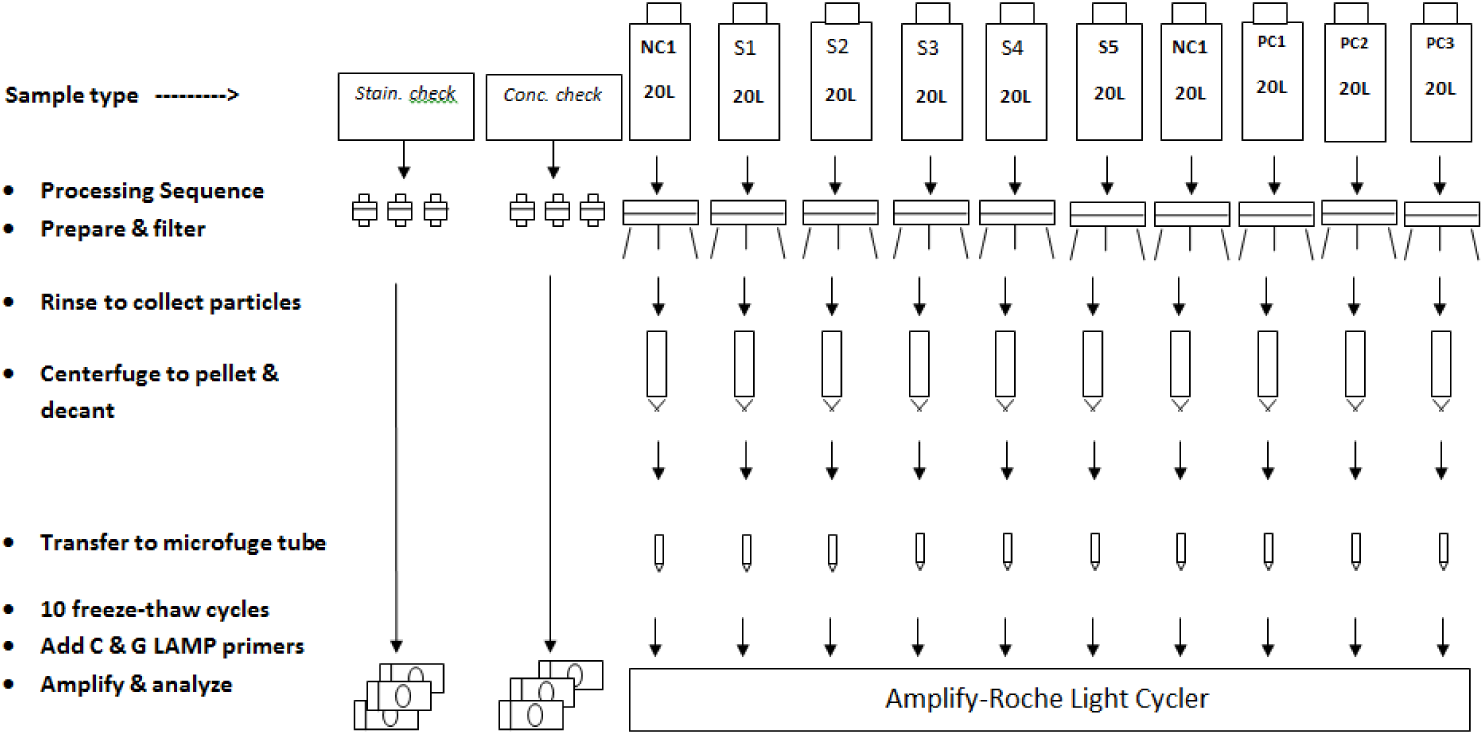
Typical water sample batch processing for LAMP detection.

## Results

Following initial LAMP runs to optimize conditions for the SAM-1 and EF1-α primers typical ladder banding patterns resulted from amplification of DNA from two calves (Lane 1, Calf C1 (CC1), Lanes 2 and 3, Calf 7056 (CC7056), Lane 4 *Cryptosporidium* NC, Lanes 5 and 6, *Giardia* Calf 7080 (GC7080), Lane 7 *Giardia* NC), Figure 2

**Figure 2.**
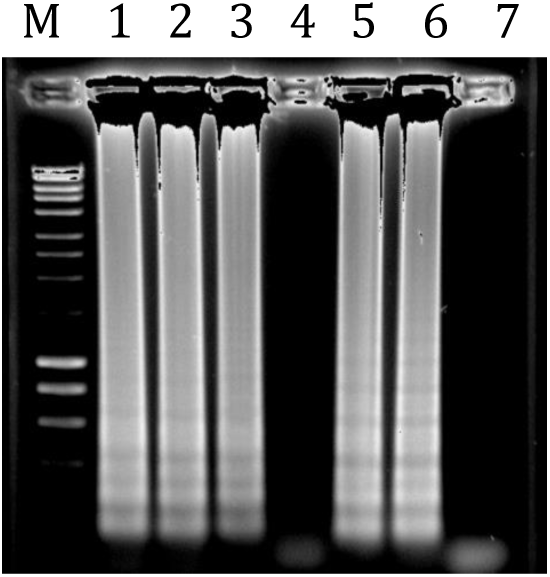
LAMP amplification of *Cryptosporidium* DNA by SAM-1 primers at 63 C for 30 minutes and *Giardia* DNA by EF1-α primers at 63 C for 60 minutes.

The sensitivity of the SAM-1 LAMP was explored by amplification of *Cryptosporidium* DNA from two sources, C1 and C7056 (CC7056) DNA, both diluted in alternate logs from 10^−1^ to 10^−10^ by LAMP using the SAM-1 primers with reaction components and amplification conditions as described above. Amplified product was observed in the resulting gel without reaching an extinction point, Figure 3.

**Figure 3.**
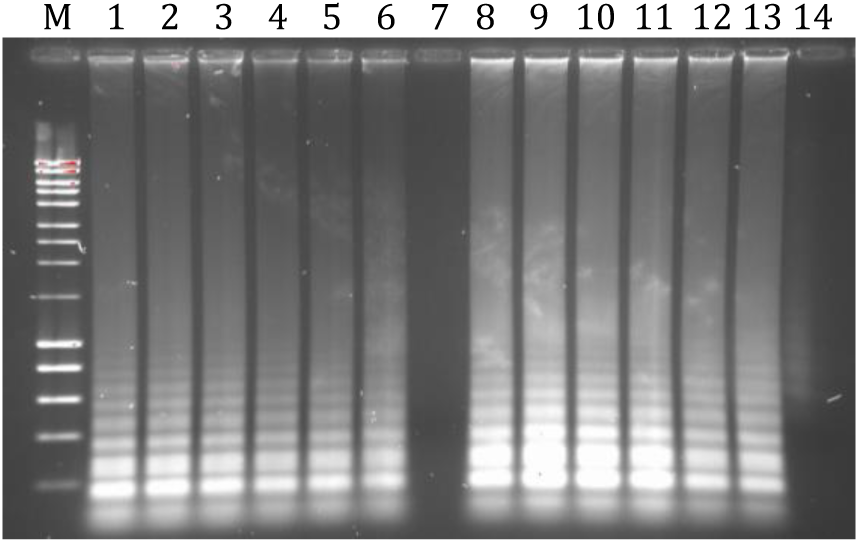
LAMP amplification of *Cryptosporidium* CC1 and CC7054 DNA diluted from 10^−1^ to 10^−10^ using SAM-1 primers at 63 C for 30 minutes, Negative controls, Lane 7, *Cryptosporidium* reagent blank; Lane 14, *Cryptosporidium* reagent blank.

The characteristic ladder banding patterns indicating amplified product for both *Cryptosporidium* DNA samples were produced at dilutions as low as tested, i.e. 10^−10^ isolated from approximately 10^7^ oocysts.

The sensitivity of the EF1-α LAMP was explored using *Giardia* DNA from two sources, calves G1 and G7080 DNA, both diluted in alternate logs from 10^−1^ to 10^−10^ by LAMP using the EF1-α primers with reaction components and amplification conditions as described above. Amplified product was observed in the resulting gel without reaching an extinction point, Figure 4. The characteristic ladder banding patterns indicating amplified product for both *Giardia* DNA samples were produced at dilutions as low as tested, i.e. 10^−10^ isolated from approximately 10^7^ oocysts.

**Figure 4.**
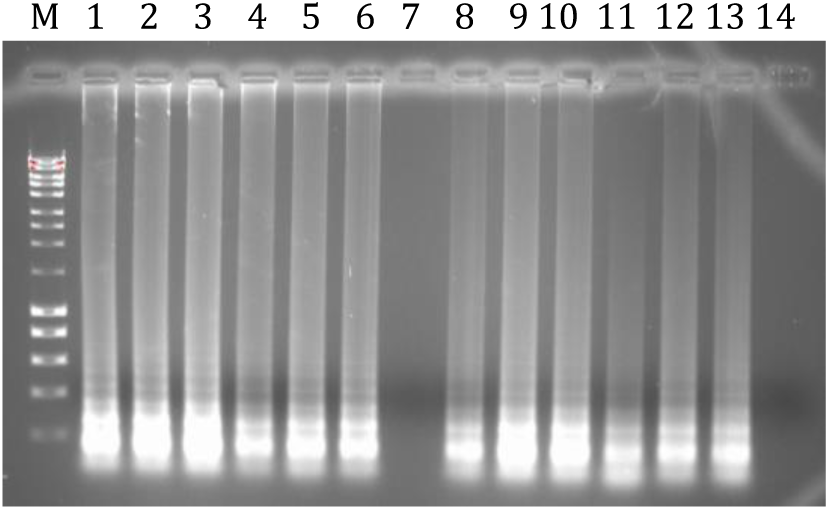
LAMP amplification of *Giardia* G1 and G7080 DNA diluted from 10^−1^ to 10^−10^ using EF1-α primers at 63 C for 30 minutes, Negative controls, Lane 7, *Giardia* reagent blank; Lane 14, *Giardia* reagent blank.

These results indicate that the SAM-1 and EF1-α LAMPs have sufficient sensitivity to amplify DNA from less than a single oocyst or cyst. Ability to amplify DNA released from oocysts and cysts without extracting and purifying the DNA was tested in duplicate pairs of 10 and 100 *Cryptosporidium* oocysts, the other 10 and 100 *Giardia* cysts, compared to duplicate C7056 DNA and duplicate G7080 DNA, with reagent blanks for *Cryptosporidium* and *Giardia*. Characteristic ladder banding was observed, Figure 5, for freeze-thawed oocysts (Lanes 1 and 2) and freeze-thawed G7080 cysts (Lanes 3 and 4) and for C7056 DNA (Lanes 5 and 6) and G7080 DNA (Lanes 7 and 8), Lanes 9 and 10 were *Cryptosporidium* and *Giardia* negative controls respectively.

**Figure 5.**
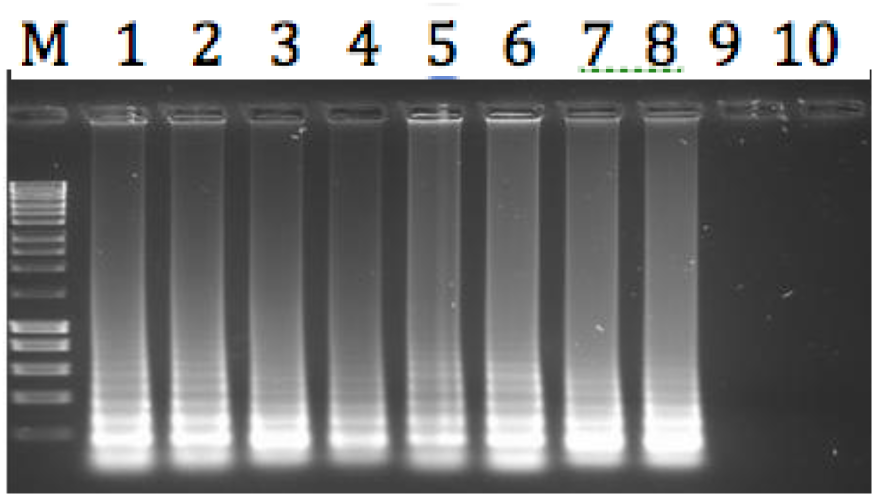
LAMP reaction components w/ 10 freeze-thaw cycles compared to C & G DNA, with negative controls.

The next step explored using the qualitative gel endpoint was to determine ability of the LAMP procedure to detect (amplify) DNA from small numbers, i.e., one, two, three, and four oocysts or cysts as would be required of a detection method suitable for application to water sample analysis. For this test, an adaptation of the drop counting procedure described previously, Ongerth & Saaed, 2013, was used to transfer single oocysts to five replicate microfuge tubes, then two oocysts to another five replicate tubes. Three and four oocysts were added each to two additional sets of four and three replicate tubes respectively. Four comparable sets of five replicates of 1 and 2, and four replicates of 3, and 3 replicates of 4 *Giardia* cysts were also prepared. Reaction components were added as described above and amplifications were completed. The *Cryptosporidium* oocyst amplifications produced amplified product for 2 of five 1-oocyst replicates, for 3 of five 2-oocyst replicates and for each of the 3 and 4 oocyst replicates, Figure 6.

**Figure 6.**
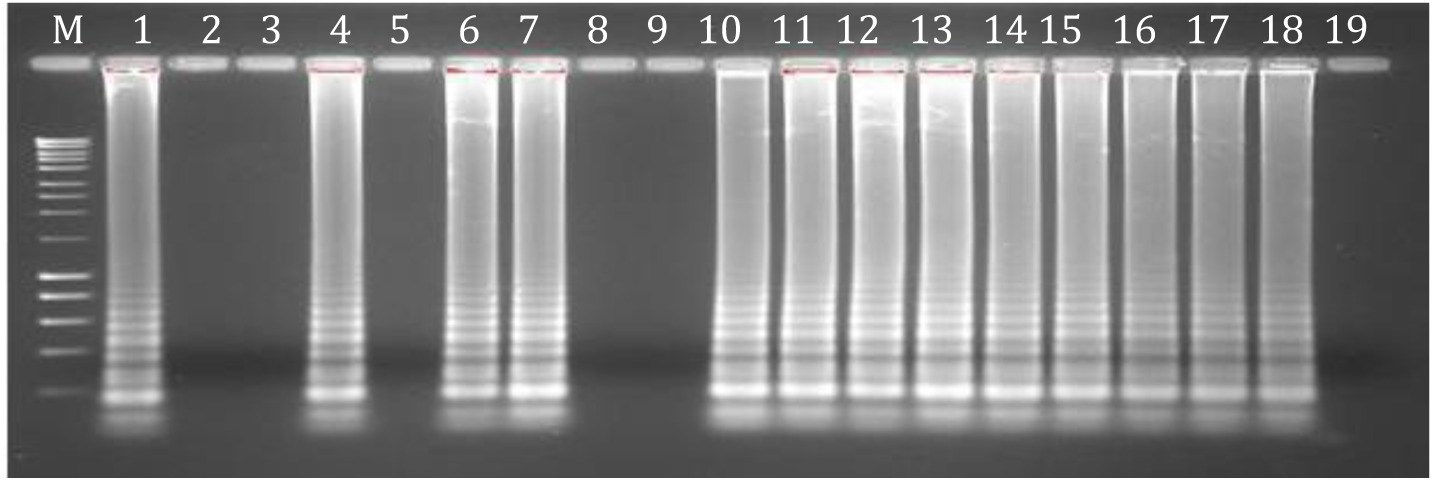
SAM-1 LAMP amplification of *Cryptosporidium* DNA from 1 oocyst (lanes 1-5), 2 oocysts (lanes 6-10), 3 oocysts (lanes 11-14) and 4 oocysts (lanes 15-17), C 7056 DNA positive (lane 18), and full reaction mix negative control (lane 19).

The *Giardia* cyst EF1-α reactions produced amplified product from 3 of 5 single cysts from 4 of 5 reactions containing 2 cysts and from each of the 4 replicated reactions each containing 3 and 4 *Giardia* cysts, Figure 7.

**Figure 7.**
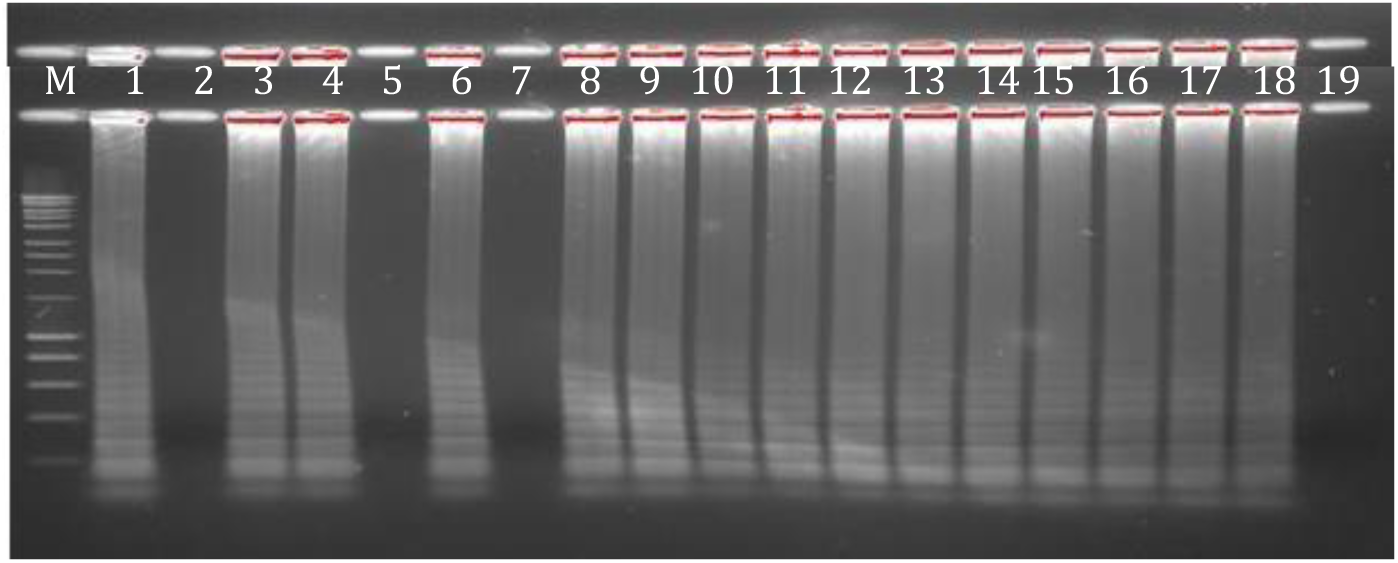
EF1-α LAMP amplification of *Giardia* DNA from 1 cyst (lanes 1-5), 2 cysts (lanes 6-10), 3 cysts (lanes11-14), 4 cysts (lanes 15-17), G 7080 DNA positive control, and full reaction mix negative control (lane 19).

The selectivity of the SAM-1 and EF1-α LAMP procedures was assessed, seeding small numbers of oocysts and cysts to actual pellet formed by filtering and centrifuging naturally occurring particles obtained from a local surface stream. The water sample was processed to form a pellet as described previously, Ongerth, 1994. Then, portions of the pellet were IMS-cleaned, then seeded with replicated numbers of oocysts and cysts ranging from 1 to 4 as described above. Replicates of both organisms included 5 each of both 1 and 2 organisms, and 4 of 3 and 3 of 4 organisms respectively. The seeded pellet volumes comprised 10.2 µL of the LAMP reactions, reaction component made up the total to 25 µL. The seeded pellets, containing all manner of naturally occurring organisms present in the stream, were subjected to the 10 freeze-thaw cycles and then amplified directly without concentrating DNA. Similar to results shown in Figure 6 and Figure 7, above, in the pellet seeded with single oocysts 2 of 5 produced amplified product; 3 of 5 seeded with 2 oocysts produced amplified product; and each of the replicates containing 3 and 4 oocysts resulted in amplified product, Figure 8.

**Figure 8.**
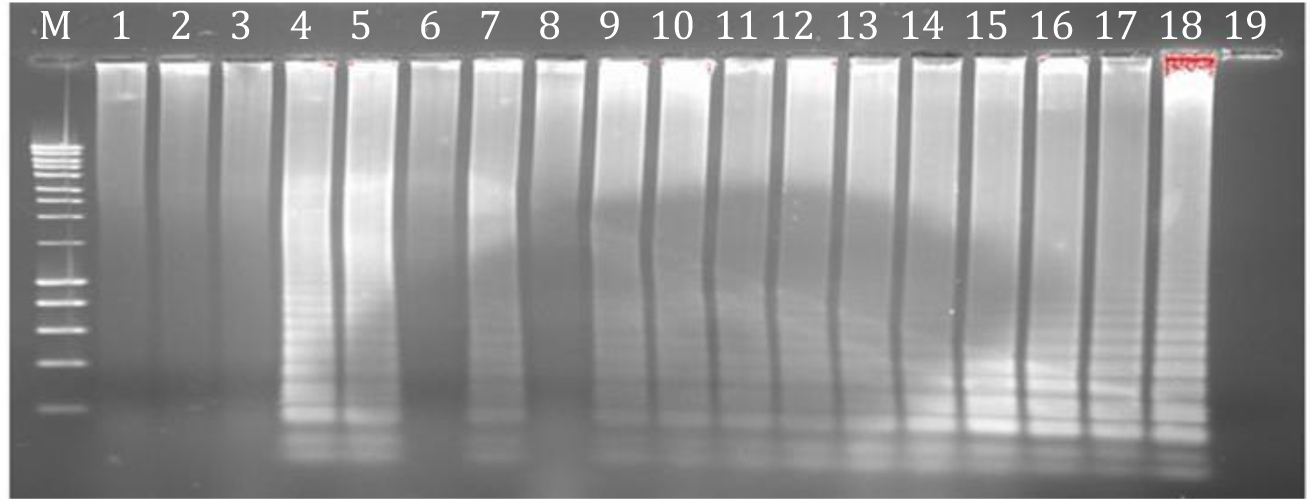
SAM-1 LAMP amplification of *Cryptosporidium* DNA from addition of oocysts to an IMS-cleaned surface water sample: 1 oocyst (lanes 1-5), 2 oocysts (lanes 6-10), 3 oocysts (lanes 11-14) and 4 oocysts (lanes 15-17), C7056 DNA positive control (lane 18), and full reaction component negative control (lane 19).

Results from amplifying pellet seeded with single *Giardia* cysts 3 of 5 resulted in amplified product; 3 of 5 seeded with 2 oocysts also resulted in amplified product; and each of the replicates containing 3 and 4 oocysts resulted in amplified product, Figure 9.

**Figure 9.**
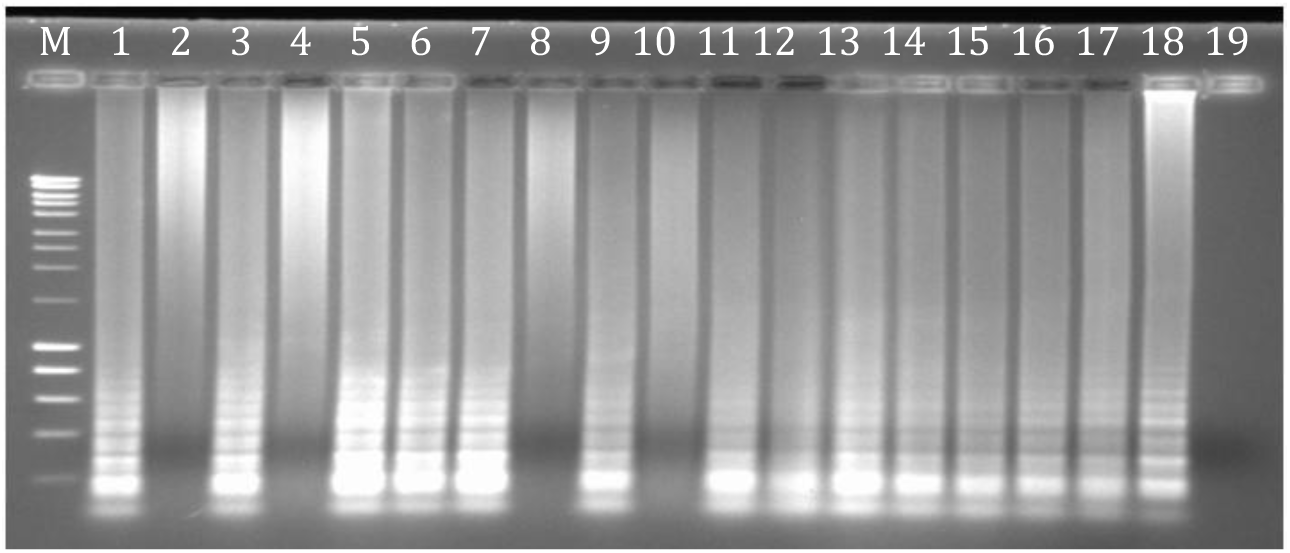
EF1-α LAMP amplification of *Giardia* DNA from addition of cysts to an IMS-cleaned surface water sample: 1 cyst (lanes 1-5), 2 cysts (lanes 6-10), 3 cysts (lanes 11-14) and 4 cysts (lanes 15-17), G7080 DNA positive control (lane 18), full reaction component negative control (lane 19).

Having observed LAMP ability to amplify DND of both organisms released from one to four oocysts and cysts by freeze-thaw cycles experiments, ability of LAMP to detect Cryptosporidium and Giardia DNA in surface water containing naturally occurring organisms was examined. Using the simplified procedure for water processing with analysis by LAMP, 5 replicate 10 L water samples with accompanying positive and negative control showed positive results for the presence of *Cryptosporidium* DNA but not *Giardia* DNA after a complete reaction time of 60-120 minutes at 63oC in LAMP assays, Figure 10 and Figure 11. Only the positive control for *Giardia* showed *Giardia* product. The LAMP assay amplified the oocyst DNA recovered from each 10 L water sample. However, only the positive control seeded with cysts resulted in amplification.

**Figure 10.**
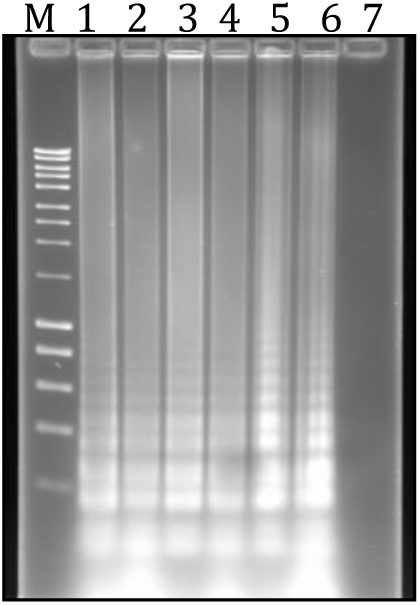
Detection of *Cryptosporidium* DNA by SAM-1 LAMP in pellets of 5 × 10 L subsamples of surface water (lanes 1-5) for comparison to IMS-based analysis, Table. Lane 6, seeded positive control, Lane 7, MilliQ water negative control.

**Figure 11.**
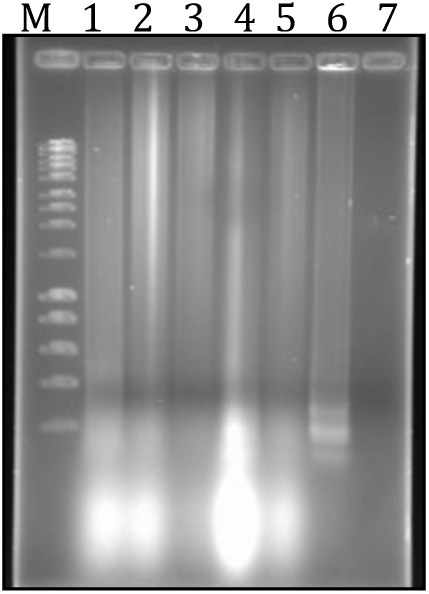
Detection of *Giardia* DNA by EF1-α LAMP in pellets of 5 × 10 L subsamples of surface water (lanes 1-5) for comparison to IMS-based analysis, Table 1. Lane 6, seeded positive control, Lane 7, MilliQ water negative control.

In the same 50 L (analyzed as 5 replicate 10 L aliquots) water sample analyzed by Method 1623, one oocyst was found on the microscope slides by immune-fluorescence microscopy (IFA) examined after monoclonal antibody stained and mounted under a cover slip for enumeration of target organisms. One of 5 replicate 10 L water samples and seeded positive control were thus shown to be positive for *Cryptosporidium* presence by the conventional IMS-IFA procedure. The conventional procedure did not detect any *Giardia* cysts in any of 5 replicate 10 L of water samples. As described above, this corresponds to results of the LAMP assay applied to freeze-thawed pellet from the simplified (filtration-centrifugation) water processing procedure

**Table 1.** Analysis of *Cryptosporidium* and *Giardia* by IMS-based procedure for comparison to detection of *Cryptosporidium* and *Giardia* in the pellet by LAMP

| Sample No. | Sampling Location | Sample Turbidity NTU | Sample Volume Litre | Crypto. No. In Sample | Giardia, No. In Sample | Crypto. Recovery % | Giardia Recovery % | Crypto Conc. No./ L | Giardia Conc. No./ L |
| --- | --- | --- | --- | --- | --- | --- | --- | --- | --- |
| 1a | UoW Ck | 0.35 | 10 | 0 | 0 | 30.7 | 87.0 | 0.0 | <0.11 |
| 1b |  |  | 10 | 0 | 0 | 30.7 | 87.0 | 0.0 | <0.11 |
| 1c |  |  | 10 | 0 | 0 | 30.7 | 87.0 | 0.0 | <0.11 |
| 1d |  |  | 10 | 1 | 0 | 30.7 | 87.0 | 0.33 | <0.11 |
| 1e |  |  | 10 | 0 | 0 | 30.7 | 87.0 | 0.0 | <0.11 |
| 1 Total |  |  | 50 | 1 | 0 | 30.7 | 87.0 | 0.07 | <0.023 |

Analysis of a 60 L water sample processed in 5 × 10 L subsamples for *Cryptosporidium* and *Giardia* by the IMS-based method, with an additional 10 L subsample seeded and processed to measure recovery efficiency (matrix effect) resulted in finding a single *Cryptosporidium* oocyst in 1 of the 5 10 L subsamples and no *Giardia* cysts in any of the 50 L, Table 1.

A companion 60 L of water collected from the same location at the same time was processed identically to the 60 L above in the filtration and centrifuging steps to form five pellets from five replicate 10 L subsamples. Pellet volumes, approximately 0.1 mL, were transferred to microfuge tubes, subjected to freeze-thaw cycles as described above, vortexed, then, 10.2 μL of pellet supernatant was processed by LAMP for amplification along with relevant positive and negative controls as described above. Reaction products indicated that *Cryptosporidium* oocysts were present in each of the five 10 L subsamples, Figure 10.

However, LAMP results showed no specific amplification corresponding to the finding of no *Giardia* cysts in any of the five 10 L subsamples, Figure 11.

The multiplex and quantitative capability of the *Cryptosporidium* plus *Giardia* LAMP was determined using real-time quantitative PCR with a fluorescence endpoint using SYTO13 and absolute endpoint determination using a Roche LightCycler 480. Using the reaction mix containing both primer sets, LAMP *Cryptosporidium* DNA using the SAM-1 gene primers and *Giardia* DNA using the EF1-α gene primers were both amplified and were distinguishable as shown with fluorescence detection in the LightCycler 480 format, Figure 12, and more clearly in LightCycler output expressed in terms of the 2^nd^ derivative melt curve analysis, Figure 13.

**Figure 12.**
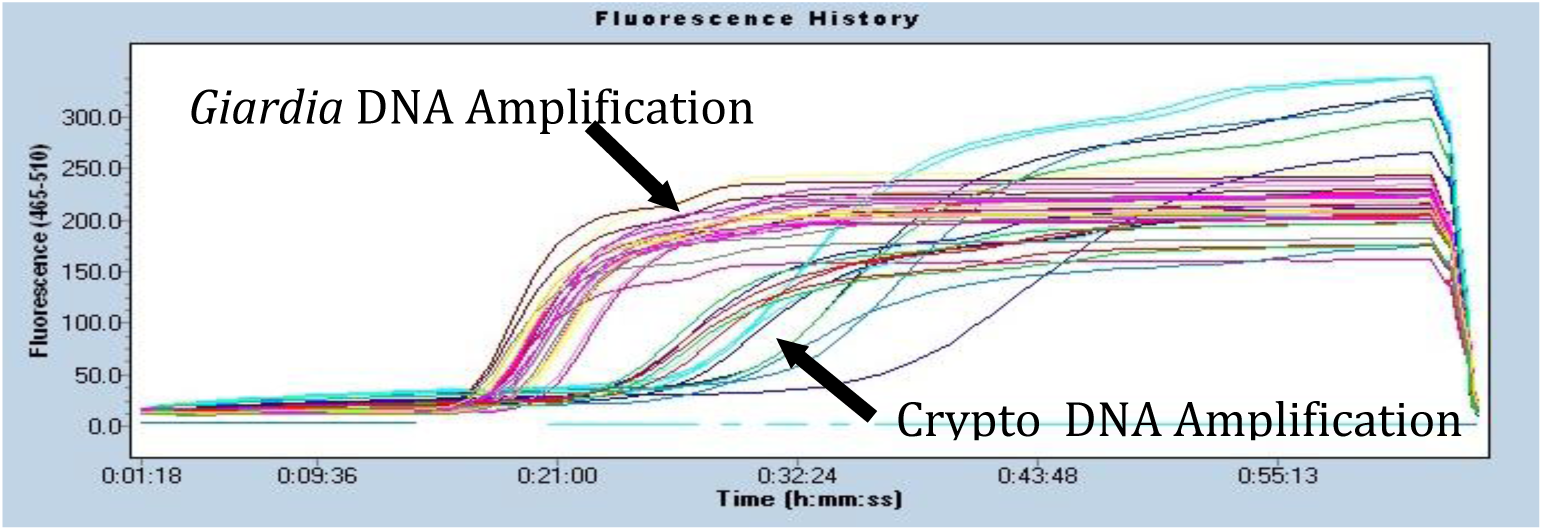
Multiplex *Cryptosporidium* & *Giardia* DNA (SAM-1 + EF1-α) Amplification vs. Time, Roche Light Cycler 480

**Figure 13.**
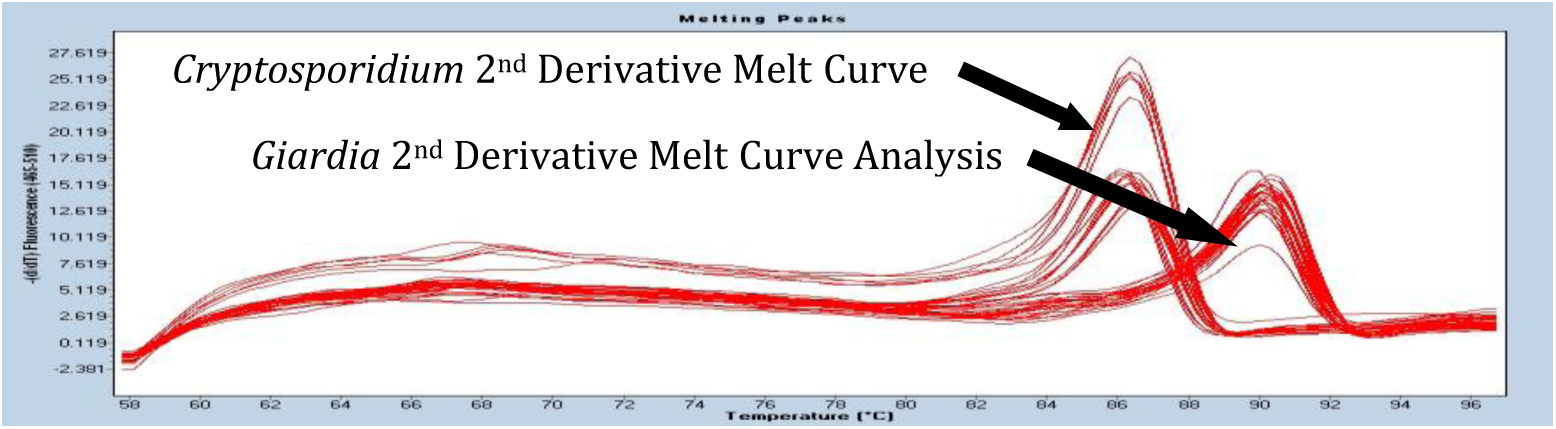
Cryptosporidium and Giardia LAMP (SAM-1 + EF1-α) product 2nd derivative melt curve Analysis

The sensitivity of the LAMP amplification for *Cryptosporidium* and *Giardia* DNA determined by dilution, was determined with fluorescence detection using the LightCycler 480. The LAMP procedure amplified the serially diluted *Cryptosporidium* and *Giardia* DNA for each dilution from the highest sample DNA concentration of 10^−8^ to as little as 10^−12^, Figure 14 and Figure 15.

**Figure 14.**
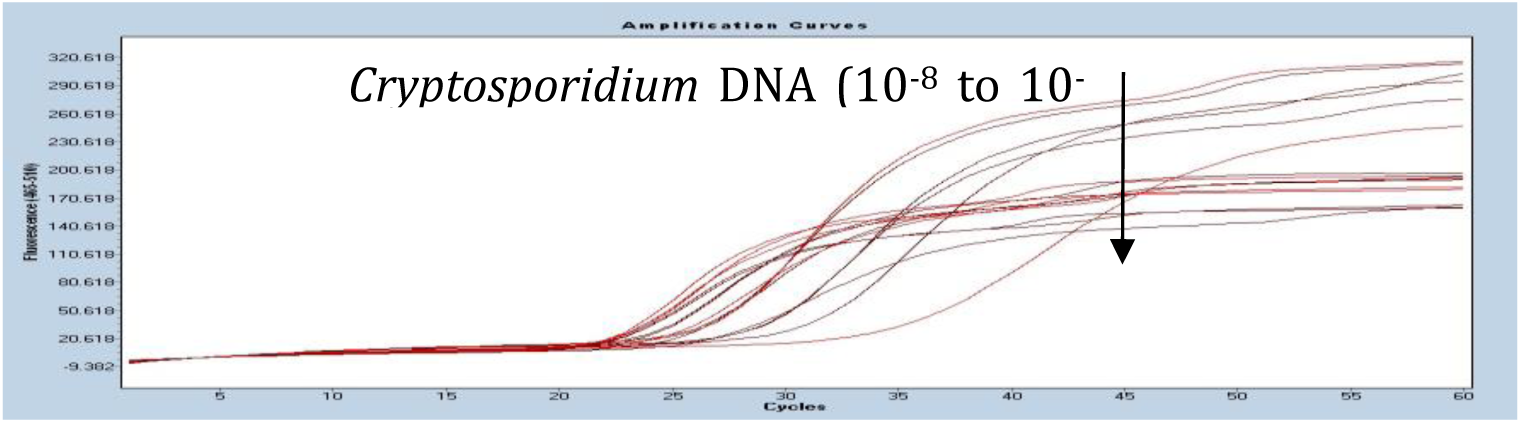
LAMP amplified *Cryptosporidium* DNA product (fluorescence) vs. Time, Roche Light Cycler 480, ½ log serially diluted DNA (10^−8^, 10^−8.5^, 10^−9^, 10^−9.5^, 10^−10^, 10^−10.5^, 10^−11^, 10^−11.5^, 10^−12^).

**Figure 15.**
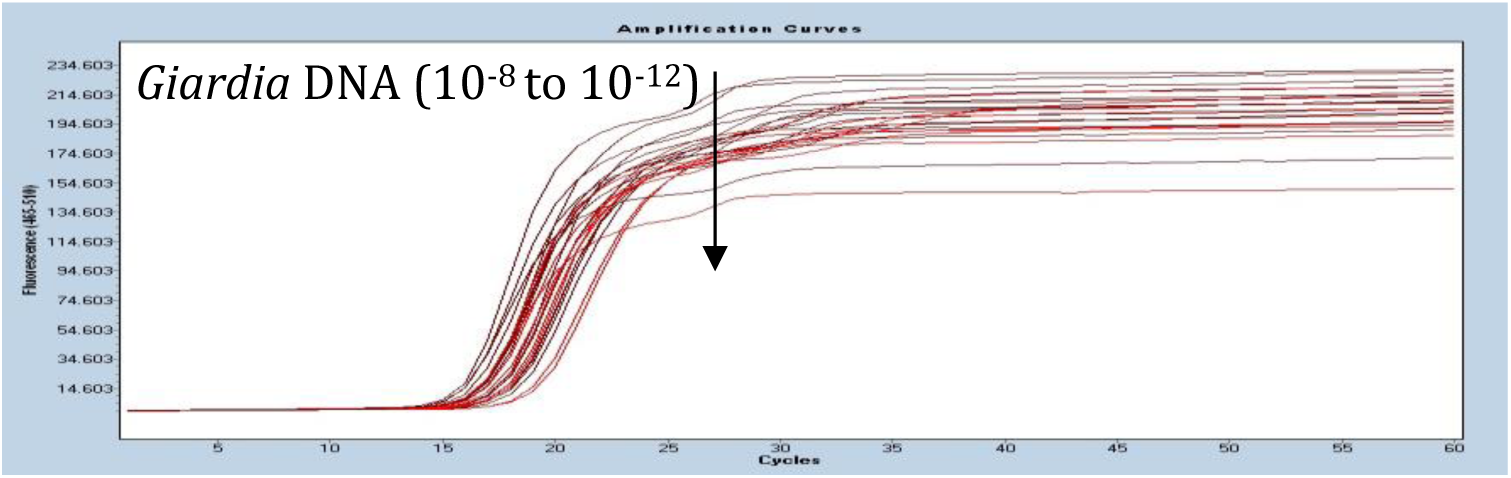
LAMP *Giardia* DNA Product (fluorescence) vs. Time. Roche Light Cycler 480, 1/2 log serially diluted (10^−8^, 10^−8.5^, 10^−9^, 10^−9.5^, 10^−10^, 10^−10.5^, 10^−11^, 10^−11.5^, 10^−12^).

The sensitivity of the LAMP amplification of *Cryptosporidium* and *Giardia* DNA in oocysts and cysts subject only to freeze-thaw cycles, was determined with fluorescence detection using the LightCycler 480 format. Amplification from of known numbers of *Cryptosporidium* oocysts and *Giardia* cysts and product (fluorescence) show the potential for discriminating between small numbers of organisms (1, 3, 7, 10, 100, 1000 and 10000) as illustrated in Figure 16 and Figure 17.

**Figure 16.**
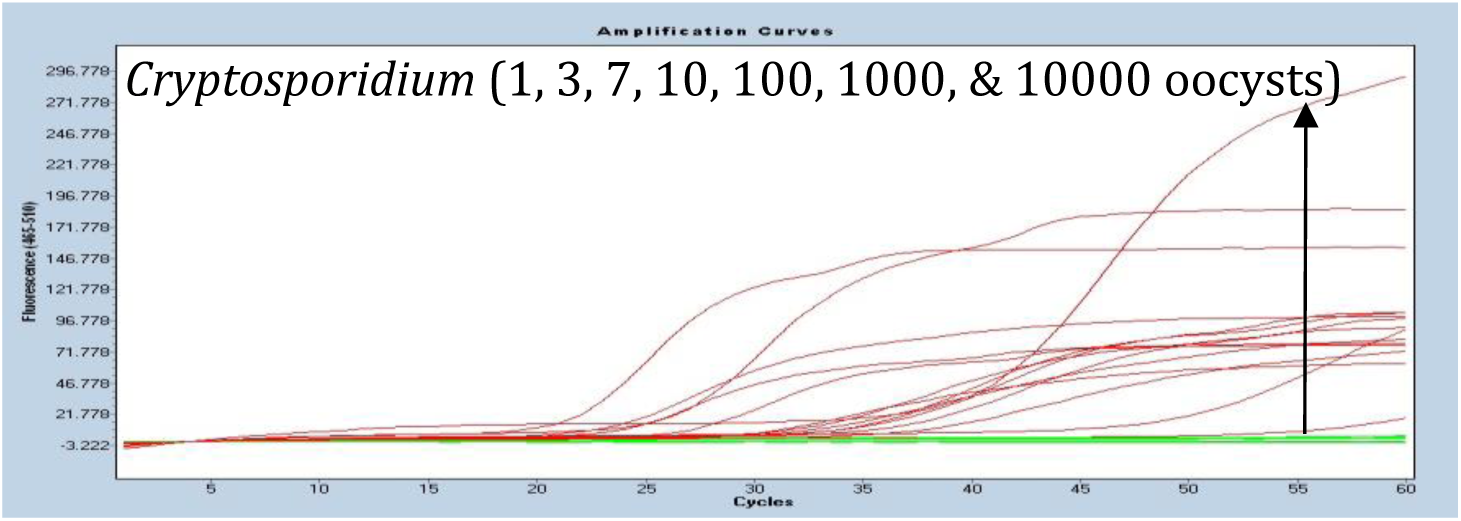
LAMP amplificaiton product DNA fluorescence from 1, 3, 7, 10, 100, 1,000 and 10,000 *Cryptosporidium* oocysts vs. Time. Roche Light Cycler 480

**Figure 17.**
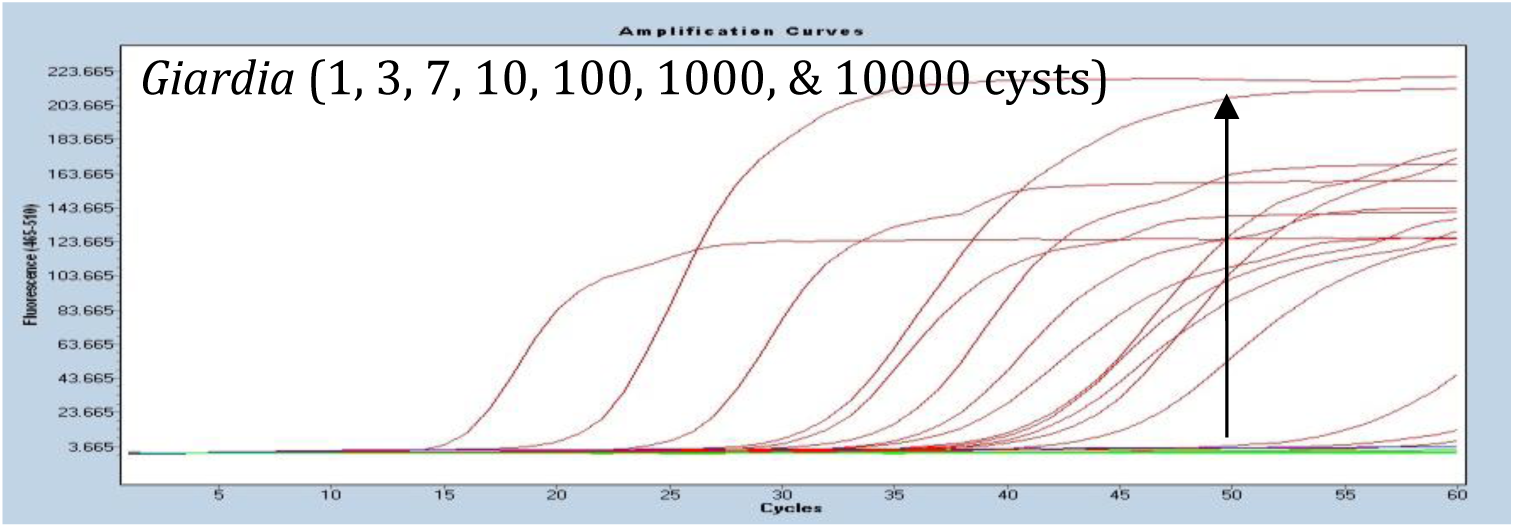
LAMP amplification product DNA fluorescence from 1, 3, 7, 10, 100, 1,000 and 10,000 *Giardia* cysts vs. Time. Roche Light Cycler 480

Application of the LAMP DNA amplification approach to the detection of *Cryptosporidium* and *Giardia* in water is not common but not novel either. As described above, previous work has described the development and application of LAMP for detection of *Cryptosporidium* (Karanis et al, 2007) and of *Giardia* (Plutzer et al, 2009) in various environmental media including surface water, drinking water, and both raw and treated sewage (Gallas-Lindeman et al., 2013, 2015). It is key features of the LAMP procedure to detect *Cryptosporidium* and *Giardia* DNA in complex media and with a high degree of sensitivity, demonstrated previously and illustrated by results presented here that are worth some discussion and make possible and have led to the simplified scheme for monitoring these pathogens in aquatic media. Repeating, for emphasis in the following discussion, the outstanding features of the LAMP amplification procedure are: 1) extreme sensitivity and 2) elegant specificity. Associated features of simplicity and low cost widely described previously (e.g., Notomi et al. 2015) contribute to factors making it attractive for environmental applications.

The sensitivity of any nucleic acid (NA) amplification procedure depends on its ability to amplify available NA copies. This feature is not always clearly elucidated in work describing, for example, PCR, qPCR, RT qPCR ability to detect *Cryptosporidium* and/or *Giardia*. Typically, the problem is framed in terms of ability to detect 1 or more oocysts or cysts without referring to the specific process by which the amplification occurs. Every amplification employs primers homologous to a region of a specific gene of *Cryptosporidium* or *Giardia*. In this work our primers made use of defined segments of the SAM-1 gene of *Cryptosporidium* and of the EF1-α gene of *Giardia*. Hence, at least a single copy of the target gene must be present in the reaction for amplification to occur assuming ideal conditions. Accordingly, critical information controlling the theoretical limit of detection is the number of copies of the relevant gene contained in a single oocyst or a single cyst. The physical morphology of an oocyst dictates that it contains 4 sporozoites each containing presumably a single copy of the SAM-1 gene. Thus, a single oocyst may contain 4 copies of this gene available for amplification. A *Giardia* cyst typically contains 2 nuclei, each of which may contain from 6-10 copies of the 5 or so chromosomes, on one of which will be located the EF1-α gene. Accordingly, a single cyst may contain as many as 2 ⨯ (6-10) or 12-20 copies of the EF1-α gene and available for amplification. Early work and recent summaries (Notomi et al, 2000; Mori et al, 2013; Notomi et al 2015) have shown LAMP to have significant advantages compared to conventional PCR and to be theoretically capable of amplifying a single copy of DNA. Numerous LAMP application reports (e.g. Karanis et al, 2007; Bakheit et al, 2007, Plutzer et al, 2009) have demonstrated LAMP applied to detection of both *Cryptosporidium* and of *Giardia* to amplify their DNA to levels diluted to that equivalent to less than a single oocyst or cyst. The application of the SAM-1 LAMP for *Cryptosporidium* and the EF1-α LAMP for *Giardia* in this work corresponds well to the previous work. In Figures 3 and 4 our application of the two LAMPs were able to detect *Cryptosporidium* DNA and *Giardia* DNA extracted from oocysts diluted to quantities equivalent to significantly less than a single organism. This sensitivity was further demonstrated in ability to detect DNA from as few as a single oocyst and cyst released for amplification by freeze thaw cycles, Figures 5, 6, and 7. The inconsistent amplification of low numbers of oocysts and cysts, i.e. 1’s and 2’s, as observed in Figures 6 and 7 and in Figures 8 and 9, we attribute to the age of the organisms used in those experiments. The organisms had been isolated approximately 6 months previous allowing for a proportion of them to have lost DNA contents preventing amplification.

The selectivity of LAMP resulting from the multiple primer sets and the related insensitivity to environmental interferences common to PCR detection schemes is a major asset. This selectivity, combined with its sensitivity was illustrated in this work by the amplification of *Cryptosporidium* and of *Giardia* DNA released from oocyst and cyst numbers including 1, 2, 3 and 4, seeded to replicates of the unseparated pellet from filtration of 10 L aliquots of a local surface stream (turbidity, 1.5 NTU), Figures 8 and 9. This was further illustrated by finding positive *Cryptosporidium* amplification directly from a pellet from filtration of an unseeded 10 L surface water sample accompanied by a separate 10 L aliquot of the same surface water processed by the conventional (e.g. USEPA Method 1623) filtration, IMS, IFA, microscopy procedure,. That procedure also found a single oocyst in its 10 L aliquot but no *Giardia*, Figures 10 and 11.

The novel and potentially very useful aspect of the results observed here is ability of LAMP to amplify DNA from target organisms, specifically *Cryptosporidium* and *Giardia*, that are collected from surface water samples of suitable volume, here 10 L, by filtration and concentrated solely by centrifuging to pellet. As has been demonstrated previously, concentration from 10-20 L to a pellet of 0.1-0.2 mL in this 2-step process is practical (Ongerth, 1994). Subjecting the decanted pellet to freeze-thaw cycles from which LAMP can be applied using 2-10 μL of the homogenized product is capable of producing potentially quantifiable results for both *Cryptosporidium* and *Giardia* in the single sample.

An important feature of the work demonstrated here is the multiplexing of the SAM-1 and EF1-α primers in a single reaction mix. Other multiplex LAMP applications have been described in the literature illustrating its practicality (e.g., Iseki et al, 2007; Nurul Najian et al, 2016; Kim et al, 2019). The ability of more than one primer set to recognize its own target and produce amplification in the presence of extraneous DNA/RNA signal is another illustration of the specificity contributing to the power and the flexibility of the procedure and its significant advantages compared to PCR, Hardinge et al., 2019.

It is clear that the success of this procedure depends on the presence of DNA for amplification so that intact oocysts and cysts that have lost cellular contents would not enumerated. Interpretation of the public health significance of such data must take this possibility into account. Determining the typical proportion of oocysts and cysts having intact as opposed to empty contents at important sampling locations, e.g. water supply intakes, would permit objective interpretation.

In light of the relatively long history of LAMP effectiveness for monitoring *Cryptosporidium* and *Giardia* in environmental media, questions regarding acceptance and broad application should be addressed. Advantages of the procedure have been cataloged above. In our experience through this project, and from discussion with others having applied LAMP (i.e. Karanis and Plutzer) disadvantages are few. Any lab currently using RT qPCR applied to microbial monitoring in water or wastewater will have the necessary expertise and understanding of essential procedural details. The high degree of sensitivity of the procedure may perhaps be considered its principal need for care. The procedure may be subject to cross-contamination or introduction of alocthanous but specific DNA or RNA if suitable care is not taken. However, lab procedures for contamination control are well within common lab practice. We experienced some difficulty in maintain “clean” negative controls in the initial period of LAMP application. But then we (engineers but experienced in work with *Cryptosporidium* and *Giardia*) were not experienced in molecular biology lab procedure, not having previous PCR lab experience. The sensitivity, specificity, and insensitivity to common PCR interferences, and simplicity of procedural and equipment requirements provide strong motivation for others to explore LAMP application potential further.

